# Phonological representations of auditory and visual speech in the occipito-temporal cortex and beyond

**DOI:** 10.1101/2024.07.25.605084

**Authors:** Alice Van Audenhaege, Stefania Mattioni, Filippo Cerpelloni, Remi Gau, Arnaud Szmalec, Olivier Collignon

## Abstract

Speech is a multisensory signal that can be extracted from the voice and the lips. Previous studies suggested that occipital and temporal regions encode both auditory and visual speech features but their precise location and nature remain unclear. We characterized brain activity using fMRI (in male and female) to functionally and individually define bilateral Fusiform Face Areas (FFA), the left Visual Word Form Area (VWFA), an audio-visual speech region in the left Superior Temporal Sulcus (lSTS) and control regions in bilateral Para-hippocampal Place Areas (PPA). In these regions, we performed multivariate patterns classification of corresponding phonemes (speech sounds) and visemes (lip movements). We observed that the VWFA and lSTS represent phonological information from both vision and sounds. The multisensory nature of phonological representations appeared selective to the anterior portion of VWFA, as we found viseme but not phoneme representation in adjacent FFA or even posterior VWFA, while PPA did not encode phonology in any modality. Interestingly, cross-modal decoding revealed aligned phonological representations across the senses in lSTS, but not in VWFA. A whole-brain cross-modal searchlight analysis additionally revealed aligned audio-visual phonological representations in bilateral pSTS and left somato-motor cortex overlapping with oro-facial articulators. Altogether, our results demonstrate that auditory and visual phonology are represented in the anterior VWFA, extending its functional coding beyond orthography. The geometries of auditory and visual representations do not align in the VWFA as they do in the STS and left somato-motor cortex, suggesting distinct multisensory representations across a distributed phonological network.

**Significance statement:** Speech is a multisensory signal that can be extracted from the voice and the lips. Which brain regions encode both visual and auditory speech representations? We show that the Visual Word Form Area (VWFA) and the left Superior Temporal Sulcus (lSTS) both process phonological information from speech sounds and lip movements. However, while the lSTS aligns these representations across the senses, the VWFA does not, indicating different encoding mechanisms. These findings extend the functional role of the VWFA beyond reading. An additional whole-brain approach reveals shared representations in bilateral superior temporal cortex and left somato-motor cortex, indicating a distributed network for multisensory phonology.

## 1. Introduction

Speech processing is fundamentally multisensory. Abstract phonological units (i.e., minimal categorical units of language) can be accessed through phonemes (speech sounds) but also through visemes, which are categories of lip movements corresponding to speech sounds (Fisher et al., 1968). Visible speech cues significantly support auditory speech perception, especially in challenging listening conditions (Sumby and Pollack, 1954) and the well-known McGurk effect illustrates the mandatory fusion that can happen in our mind between phonemes and visemes (McGurk and MacDonald, 1976).

Which brain regions support the joint representation of visemes and phonemes? Posterior superior temporal regions are often considered as the core phonological hub in auditory speech perception (Hickok and Poeppel, 2007; Chang et al., 2010; Bhaya-Grossman and Chang, 2022). Beyond auditory phoneme representation, studies on audio-visual speech processing (Capek et al., 2008; Okada and Hickok, 2009; Stevenson et al., 2010; Nath et al., 2011) or even visual speech processing only (Calvert et al., 1997; Paulesu et al., 2003; Hall et al., 2005; Bernstein et al., 2011) have reported activations in pSTG or pSTS (see Venezia et al. (2018) for a meta-analysis).

Beyond the pSTS, it was more recently suggested that the ventral occipito-temporal cortex (VOTC) may be involved not only in visual but also in auditory phonological processing (Hauswald et al., 2018; Nidiffer et al., 2023). Activations in the VOTC have been consistently reported in studies involving lipreading and audiovisual speech (Campbell et al., 2001; Paulesu et al., 2003; Hauswald et al., 2018; Nidiffer et al., 2023). Most authors relate the activation in the VOTC for lipreading to the fusiform face area (FFA), but this assumption was never tested directly (Bernstein et al., 2014). Another region of interest that has received comparatively lower attention in this context is the visual word form area (VWFA). This brain region located in the left VOTC is classically defined as the core of orthographic processing in reading (Cohen et al., 2002; Dehaene et al., 2005; Vinckier et al., 2007). Its sensitivity to written words is traditionally believed to develop from a preference for low-level visual features that match those of the orthographic code, such as a foveal bias (Hasson et al., 2002) or a preference for specific shapes, line-junctions or spatial frequencies (Vogel et al., 2012; Szwed et al., 2011). Yet, an alternative hypothesis for the functional specialization of the VWFA posits that this region serves as an interactive system between the visual and language network (Price and Devlin, 2011; Qin et al., 2021). This region indeed exhibits strong anatomical and functional connections with the language system, even before reading acquisition occurs (Saygin et al., 2016). In line with this idea, there is growing evidence for spoken-language activations and phonological processing in VWFA (Zhao et al., 2017; Taylor et al., 2019; Pattamadilok et al., 2019), even in the absence of written material (Yoncheva et al., 2010; Planton et al., 2019; Qin et al., 2021; Wang et al., 2022). Given that both reading and lipreading require audio-visual matching of visual units (graphemes or visemes) to phonemes, the VWFA, and particularly the anterior section of the VWFA (Lerma-Usabiaga et al., 2018) emerges as a good candidate region for supporting auditory and visual phonological representations and their integration in the VOTC.

In our study, we first individually and functionally identified regions of interest (ROIs) within the occipital and temporal cortices for faces (FFA), words (VWFA), multisensory speech (lSTS) and places (parahippocampal area, or PPA) as a control region in VOTC. Using multivariate pattern (MVP) decoding of phonological units presented in audition and vision, we aimed to test which of these ROIs selectively support phoneme and viseme processing. Second, using cross-modal decoding, we investigate whether putative regions housing both auditory and visual phonological representations present a similar activation geometry across modalities, suggesting a partially shared phonological coding across the senses.

## 2. Materials and methods

### 2.1 Participants

Twenty-four adults completed the fMRI study protocol (13 males, mean age ± SD = 24.04 ± 3.25, age range = 19-32 years). All participants were native French speakers and had normal or corrected-to-normal vision (corrected in the scanner with contact lenses or MRI-compatible glasses) and self-reported normal audition. Twenty-one of the 24 participants were right-handed. Participants had no reported history of dyslexia, attention disorder, psychiatric or neurological problems and were not under medical treatment for any of those problems. All participants were naive to the purpose of the experiment. They gave their informed consent before the experiment and received monetary compensation for their participation. The experimental protocol was approved by the Ethical Committee from the University Hospital of Saint-Luc (protocol n° CE-2016/23Mai/228). One subject was excluded from analyses due to excessive head motion, resulting in a total sample of 23 participants. We did not conduct formal power analysis as there was no previous study investigating the main effect of interest of this study (cross-modal decoding of phonemes and visemes). Our sample size was determined based on similar fMRI MVP studies on auditory phonological representations (Bonte et al., 2014; Correia et al., 2015; Arsenault and Buchsbaum, 2015) and on available resources.

### 2.2 Experimental design

Each participant underwent two experimental sessions for about 4 hours of testing in total. The first session included a brief training outside the scanner to familiarize participants with visual syllables, followed by the MRI acquisition of the structural images, as well as functional scans for one of the localizer experiments and for the event-related phonological experiment (up to 10 auditory runs and 10 visual runs). In the second session, participants completed functional runs for the remaining two localizer experiments and the second half of the event-related phonological experiment (up to 10 auditory and 10 visual runs). The administration order of the localizer experiments across the two sessions was counterbalanced across participants. The visual and auditory runs of the event-related phonological experiment were alternating, and the order of presentation was counterbalanced across participants: half of the subjects started with an auditory run and the other half started with a visual run. The time inside the MRI scanner was limited to 90 minutes per session. Visual stimuli were presented on a screen through a mirror at a distance of 170 cm and with a resolution of 1920×1200 pixels. Auditory stimulation was delivered through in-ear MRI-compatible earphones (Sensimetric S14, SensiMetrics, USA) at a loud but comfortable volume, defined with each participant before the start of the MRI acquisition.

#### 2.2.1 Word-Face-Scene localizer

A visual localizer was implemented to define within a single experiment our regions of interest in the VOTC, namely the VWFA, bilateral FFA, and bilateral PPA as a control region. Images of written words, houses and faces (24 exemplars per category) were used in a block design. The written words were abstract French words in various fonts on a white background. The images of houses were collected from Internet. The faces were neutral pictures of male and female taken from the *Face Research Lab London Set* (DeBruyne & Jones, 2017). All images were made 400 x 400 pixels large, were set to black and white and equated across categories for spatial frequencies and mean luminance using the SHINE toolbox (Willenbockel et al., 2010). The stimuli sustained 4.8 degrees of visual angle on the MRI screen.

We presented 10 blocks of each visual category, resulting in 30 blocks for the total localizer experiment. In each block, 12 stimuli from one category are presented in random order, for a duration of 750ms and are followed by a 250ms fixation cross in the center of the screen (ISI). Blocks alternated between faces, houses and words conditions in a random order and were separated from each other by a 6-seconds inter-block interval (IBI). Participants performed a one-back task during the presentation of the stimuli. They were asked to press a button with their right index when an image was repeated two times in a row. Each block contained 0, 1 or 2 targets (0 to 8% of the trials). The design is illustrated in Fig. 1A.

**Figure 1.**
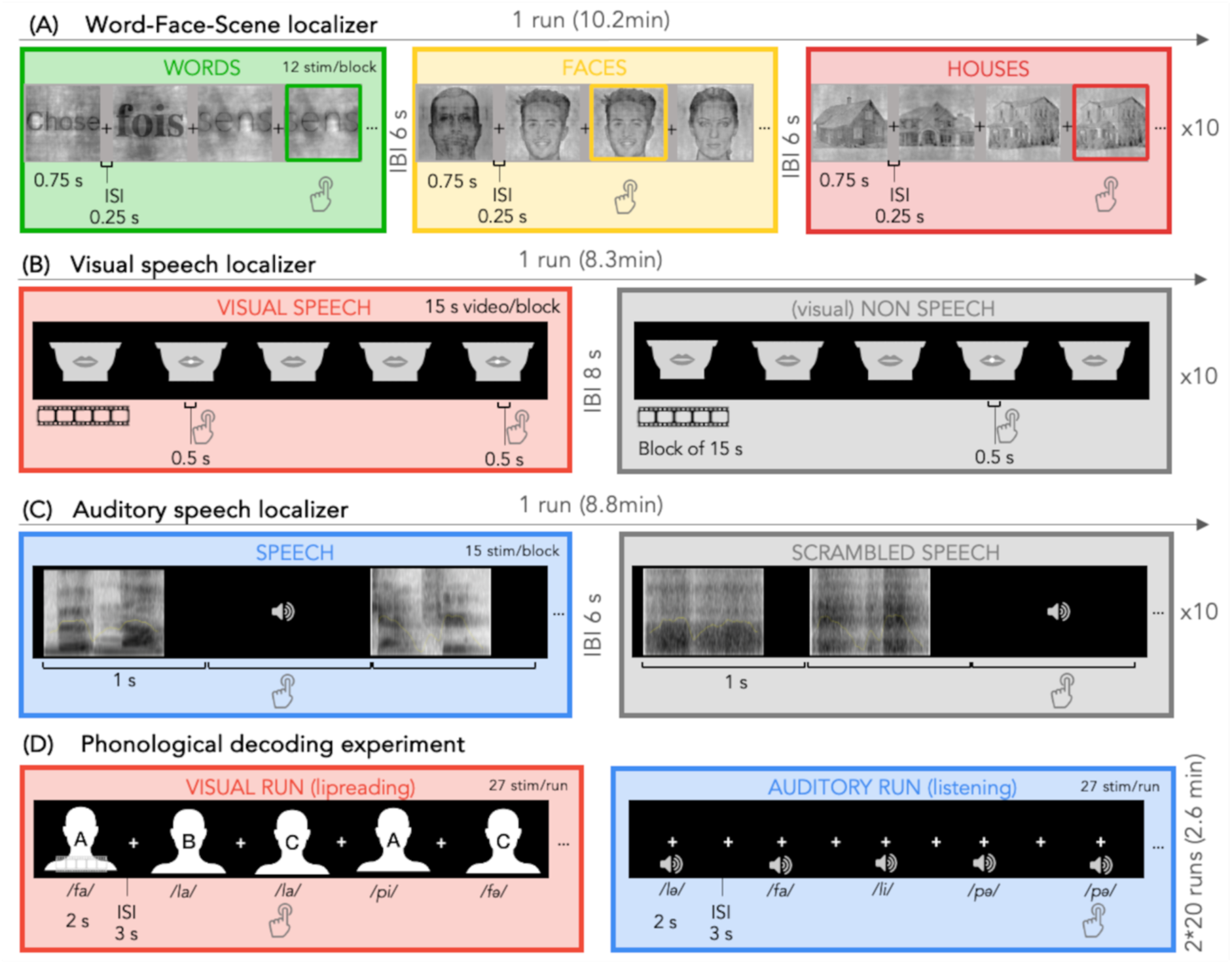
Experimental design and example of the stimuli for each of the 4 fMRI experiments. (A) Example of blocks in the 3 conditions (words, faces and houses) presented in the VOTC-localizer. Each condition was repeated 10 times. Participants performed a one-back task. (B) Example of blocks in the 2 conditions (visual speech and non-speech) of the Visual-speech localizer. Each condition was repeated 10 times. Participants were required to press when a dot appeared in the centre of the video. (C) Example of stimuli and block-design in the 2 conditions (speech and scrambled speech) of the Auditory-speech localizer. Each condition was repeated 10 times. Participants were asked to press when a target tone was heard. (D) Example of a visual and an auditory run from the phonological decoding experiment. Participants did a maximum of 20 runs in each condition. The task was to detect a repeated syllable, independently of the actor.

#### 2.2.2 Visual-Speech Localizer

Based on the findings of Bernstein and colleagues (2011), we developed a localizer experiment that would enable us to individually identify a region sensitive to visual speech in the left posterior temporal cortex, also named the temporal visual speech area (TVSA). We contrasted silent video clips of an actor speaking and clips of the same actor making non-linguistic lip movements (e.g. kiss, gurns, left and right extension, …), as illustrated in Fig. 1B. The videos were cut and masked, in order to keep only the lower part of the face up to the nose, using Hitfilm Express 2021.3 (FXhome Limited, Dolby Laboratories). The videos were presented to the participants on a black background, sustaining 8 to 10 degrees of visual angle at the participant’s distance from the screen. Participants were asked to look attentively to the video clips in both conditions and to detect when a white dot appeared in the center of the screen, in the lip area. The time of apparition of the target dots was randomly defined, but with the same number and same moment of apparition in both conditions (0, 1 or 2 targets in each block). One video clip of 15 seconds constituted one block. We presented 20 blocks in total (10 per condition), with an IBI of 8 seconds (white fixation cross) and a 10-second fixation-cross at the start and at the end of the run.

#### 2.2.3 Auditory-Speech localizer

To localize auditory regions that respond preferentially to speech and phonological processing, we implemented a block-designed auditory localizer, similarly to our Visual-Speech localizer. We presented meaningless bisyllabic pseudo-words and their scrambled version. The scrambled version control for the rhythm (speech envelope) and spectro-temporal content of the speech signal (Calce et al., 2024). By contrasting phonologically relevant speech sounds with phonologically irrelevant transformed sounds, we define broad auditory areas responsive to voice, speech and phonological processing. A large set of consonant-vowel syllables from 2 male and 2 female native French speakers were recorded in a sound-proof environment, using a digital stereo recorder at 44.1kHz. The soundtracks were concatenated using the software Praat (version 6.2.03; Boersma & Weenink, 2021; http://www.fon.hum.uva.nl/praat/) to create meaningless bisyllabic pseudo-words. To create an unintelligible version of the sounds that contained no phonological information, the same recordings were transformed using a method described in Calce et al. (2024), by applying a fast Fourier transformation (FFT) to the sounds and shuffling the Fourier components between each other within frequency bins of 200Hz. An inverse FFT was then performed on this signal and the original sound envelope was reapplied to the resulting scrambled sound. This method allows to get unintelligible vocal sounds that have similar frequency content and spectral-temporal structure to the one of the original sounds. Finally, all soundtracks - original and scrambled - were root mean square (RMS)-equalized.

The localizer consisted of one run of 20 blocks (10 per condition). In each block, 15 bisyllabic pseudo-words or their scrambled equivalent are presented (Fig. 1C). Sound clips have various durations ranging from 0.6 to 0.9s (mean duration = 0.77s) and are followed by an interstimulus interval (ISI) to get to a presentation rate of 1s per trial (mean ISI duration = 0.23s). Each block therefore had a duration of 15s. Blocks alternated between speech and scrambled conditions and were separated from each other with an inter-block interval (IBI) of 6s. The order of presentation of the blocks was randomized across participants. The localizer started and ended with an 8-second fixation cross. Participants were asked to detect a target sound (pure tone at 250Hz) in the sound sequence by pressing on a button with their right index. The number of targets (ranging from 0 to 2 in each block) was randomized and balanced across conditions. To minimize eye movements, the participants were instructed to keep their eyes open and to fixate a cross in the center of the screen during the whole duration of the task.

#### 2.2.4 Event-related phonological experiment

Syllables were presented to the participants in auditory (speech) and visual (lipreading) format in an event-related design in this fMRI experiment. Three consonants (/p/, /f/, /l/), combined with 3 vowels (/a/, /ə/, /i/) in a consonant-vowel syllable structure and pronounced by 3 different speakers (2 males) were presented, for a total of 27 unique muted videos (visual condition) and 27 soundtracks (auditory conditions). We decided to combine the consonants with several vowels and pronounced by several actors, to create visual and auditory variability that would be colinear with the feature of interest (the consonant), ensuring that the decoding is performed on the abstract representation of the phonemes, and not on low-level features of the stimuli. Consonants have been selected as feature of interest since they have more specific articulatory patterns (specific lip movements, classified into viseme categories) while vowels are classified across mouth aperture and roundness but not a specific movement. A pilot behavioral experiment involving 12 participants who did not participate in the MRI sessions guided the selection of highly discriminable syllables and ensured good performance in the auditory and lipreading task. The selected consonants (/p/, /f/, /l/) are articulated in different articulation places (bilabial, labiodental, alveolar) and manner (stop, fricative and liquid respectively), resulting in highly distinct lip movements and acoustic features (Fig 1E). The selected vowels /a/, /ə/ and /i/ are produced with various degrees of mouth aperture (open, neutral and closed respectively), and are typically articulated in the front or center of the mouth to ensure better visibility compared to more posterior vowels (e.g., /o/, /u/).

The video- and audio-clips lasted for 2 seconds and consisted of one actor repeating 3 times the same syllable. Stimuli were always followed by a 3-seconds fixation cross. The fixation cross was also present during presentation of the auditory stimuli and participants were instructed to fixate the cross during auditory runs. The experiment was divided in short runs of auditory or visual stimuli, as described in Fig. 1D. One run encompassed all stimuli in one modality, presented in random order avoiding repetitions of the same syllable except for targets. Participants were instructed to press a button with their right index when a syllable was presented 2 times in a row (one-back task), independently of the speaker’s identity. Targets appeared 2 or 3 times per run (5% of the trials) and were discarded from the subsequent analyses. Video clips were recorded using a full HD camera in MP4 format and a resolution of 1920×1080 pixels and with professional lightening set-up. They were cut, reframed, and masked using Hitfilm Express 2021.3 (FXhome Limited, Dolby Laboratories). They were presented on a black background on the screen at a rate of 25 fps and with a size of 1080 pixels wide and 1152 pixels high, sustaining 8 to 10 degrees of visual angle at the participant’s distance from the screen. Audio clips were recorded in the same set-up as previously described (see Auditory-Speech localizer) and were edited and converted into mono recordings using Praat (Boersma & Weenink, 2021) and then amplitude-normalized using the root mean square (RMS) method. To ensure optimal recording conditions for both auditory and visual modalities, the sound and videos were recorded separately and were never presented simultaneously.

### 2.3 Acquisition parameters

Participants were scanned at the University Hospital of Saint-Luc, Brussels (Belgium) with a 3 Tesla GE MRI scanner (SIGNA Premier system, General Electric Company, USA, serial number: 000000210036MR03, software version: 29LXRX29.1_R02_2131.a) with a 48-channel head coil. In the first scanning session, whole-brain T1-weighted structural MRI data were acquired (3D-MPRAGE; 1.0 x 1.0 mm in-plane resolution; slice thickness 1mm; no gap; inversion time = 0.9 s; repetition time (TR) = 2.18816 s; echo time (TE) = 0.00296 s; flip angle (FA) = 8°; 156 slices; field of view (FOV) = 256 x 256 mm; matrix size= 256 X 256). Functional images for localizers and event-related phonological experiment were T2*-weighted scans acquired with Echo Planar and gradient recalled (EP/GR) imaging (2.6 x 2.6 mm in-plane resolution; slice thickness = 2.6mm; no gap; Multiband factor = 2; TR = 1.750 s, TE = 0.03 s, 58 interleaved ascending slices; FA = 75°; FOV = 220 x 220 mm; matrix size= 84 X 84). One run of bold fMRI data was collected for the Word-Face-Scene localizer task (duration = 10.2 minutes, during which 348 volumes were acquired), one run for the Visual-Speech localizer (duration = 8.3 minutes, during which 286 volumes were acquired) and one run for the Auditory-Speech localizer (duration = 8.8 minutes, during which 302 volumes were acquired). For the event-related phonological experiment, we collected 20 visual runs and 20 auditory runs (run duration = 2.6 minutes, during which 89 volumes were acquired) for all the subjects. In some subjects, we had to exclude a few runs from the analyses, due to excessive movements in these runs. Our final data set comprised 16 subjects with 20 runs, 5 subjects with 19 runs, 1 subject with 18 runs and 1 subject with 15 runs of each modality.

### 2.4 Analyses

#### 2.4.1 Preprocessing

The MRI data were pre-processed with bidspm (v3.0.0; https://github.com/cpp-lln-lab/bidspm; Gau et al., 2022) using Statistical Parametric Mapping (SPM12 – version 7771; Wellcome Center for Neuroimaging, London, UK; https://www.fil.ion.ucl.ac.uk/spm; RRID:SCR_007037) using MATLAB 9.3.0.713579 (R2017b) on a Mac OS X computer (10.15.7). The preprocessing of the functional images was performed in the following order: slice timing correction, realignment and unwarping, segmentation and skullstripping, normalization to MNI space, smoothing. Slice timing correction was performed taking the 29th (middle) slice as a reference (interpolation: sinc interpolation). Functional scans from each participant were realigned and unwarped using the mean image as a reference (SPM single pass; number of degrees of freedom: 6; cost function: least square) (Friston et al., 1995). The anatomical image was bias field corrected, segmented and normalized to MNI space (target space: IXI549Space; target resolution: 1 mm; interpolation: 4th degree b-spline) using a unified segmentation. The tissue probability maps generated by the segmentation were used to skullstrip the bias corrected image removing any voxel with p(gray matter) + p(white matter) + p(CSF) > 0.75. The mean functional image obtained from realignement was co-registered to the bias corrected anatomical image (number of degrees of freedom: 6; cost function: normalized mutual information) (Friston et al., 1995). The transformation matrix from this coregistration was applied to all the functional images. The deformation field obtained from the segmentation was applied to all the functional images (target space: IXI549Space; target resolution: equal to that used at acquisition; interpolation: 4th degree b-spline). Preprocessed functional images were spatially smoothed using a 3D gaussian kernel (FWHM of 6 mm for the functional localizers, 2mm for the runs of the event-related phonological experiment). This method section was automatically generated using bidspm (Gau et al., 2022) and octache (https://github.com/Remi-Gau/Octache).

#### 2.4.2 General Linear Model

All pre-processed functional images were then analyzed at the subject level. We performed a mass univariate analysis with a linear regression at each voxel of the brain, using generalized least squares with a global FAST model to account for temporal auto-correlation (Corbin et al., 2018) and a drift fit with discrete cosine transform basis (128 seconds cut-off). Image intensity scaling was done run-wise before statistical modeling such that the mean image would have a mean intracerebral intensity of 100. fMRI localizer experiments were modelled in a block-design, with 3 regressors (faces, houses and words) for the Word-Face-Scene localizer and 2 regressors for the Visual-Speech localizer (visual-speech and non-speech) and the Auditory-Speech localizer (spoken and scrambled pseuso-words). The main event-related phonological experiment was modelled with three regressors corresponding to the three consonants entered into the run-specific design matrix. The onsets were convolved with SPM canonical hemodynamic response function (HRF). Nuisance covariates included the 6 realignment parameters to account for residual motion artefacts, one regressor of no-interest for the target stimuli. High motion time points were censored from the GLM if they were flagged as outliers in a run-wise manner using a robust outlier estimation algorithm. The GLM outputs for the event-related phonological experiment were concatenated into 4D maps for later multivariate pattern analysis.

#### 2.4.3 Behavioral responses during fMRI scans

We calculated a d’ (d-prime) value as an index of the sensitivity of the task in our participants. This index considers correct responses (correctly detected targets) and errors (missed targets or false alarm press). We used one value for all the runs in one modality. D’ values were calculated for each subject, in auditory and visual modality separately, using this formula: d’ = z(correct detection rate) – z(false alarm rate) (Stanislaw and Todorov, 1999). We then performed a one-sample t-test on the d’ values, to compare the performances in auditory and visual modality. Due to a technical issue, we could not record button presses for one session in one subject. The analysis of the behavioral responses thus includes 22 participants.

#### 2.4.4 VOTC-ROIs definition

Discrete regions of VOTC differ from their neighbors in their function, architectonics, connectivity, and/or internal topography (FACT; Felleman and Van Essen, 1991). We aimed to identify 5 functional regions in the VOTC: the word-selective region (VWFA), the place-selective regions (left and right PPA) and the face-selective regions (left and right FFA). To create individual ROI masks in the ventral occipito-temporal cortex, we first identified significant activation clusters at the group level for the condition of interest > control conditions, at a cluster-level threshold of p<0.05 Family Wise Error (FWE) corrected for PPA or p<0.001 uncorrected for VWFA and FFA. We then extracted the individual peak coordinates within the group-constrained cluster of interest. Starting from the individual peak coordinate within each group-constrained ROI, we drew an expanding sphere up to a radius of 10 mm (in order to include more than 200 voxels). We limited the expansion of the sphere to the boundaries of a metanalysis mask created in neurosynth using functional terms as input function (“face” for FFA, “place” for PPA and “visual word” for VWFA). This procedure ensures that ROIs are tailored to each individual functional organization while staying within reasonable distance from the known spatial location of the functionally selective regions. In line with the well-known right hemispheric dominance for face processing (Rossion et al., 2003), our localizer did not show a reliable activation peak in the left FFA by contrasting the activity for [faces] > [houses and words], in most of the subjects and at the group level (Fig 2.A). To foreshadow our results, we obtained significant auditory and visual decoding in left VWFA but not in right FFA. To ensure that the multisensory nature of VWFA was truly selective to this region (VWFA) and not widespread in the left fusiform gyrus in general (including FFA), we included a face ROI in the left hemisphere. To do so, we created a group-defined ROI by drawing a binary mask of 14mm (ensuring to include enough voxels in the later analytical steps) within the left fusiform gyrus, using precise coordinates from the literature (Fox et al., 2009). Since the left FFA and VWFA are neighboring regions, we noticed an overlap between the left FFA mask and the individual VWFA masks in most subjects. Consequently, we excluded from the group left FFA-mask all the voxels that were already encompassed by any individual VWFA mask (Fig. 2), resulting in a binary mask of 306 voxels, positioned inferiorly and anteriorly to the individual VWFA masks. In conclusion, a total of 92 individual binary masks (i.e., left VWFA, right FFA, left and right PPA across 23 subjects) and 1 group mask (left FFA) were created in the VOTC. A feature selection procedure ensured the selection of a similar number of voxels across ROIs for the decoding procedure (see section 2.4.6). Given the growing body of evidence for a neuro-functional segregation between anterior and posterior sections of the VWFA (see for example Bouhali et al., 2014; Lerma-Usabiaga et al., 2018; Nobre et al., 1994), we used “FG1” (left anterior fusiform gyrus, hereafter VWFA-ant) and “FG2” (left posterior fusiform gyrus, hereafter VWFA-post) masks from the JuBrain Anatomy Toolbox (Eickhoff et al., 2005; Amunts et al., 2020) to delineate VWFA-ant and -post at the group-level (Fig. 3). We did not run a functional localizer for VWFA-ant and VWFA-post because this complementary analysis was not included in the initial study design. To ensure functional specificity, we excluded voxels from VWFA-ant that overlapped with the group-defined left-FFA mask mentioned above.

**Figure 2.**
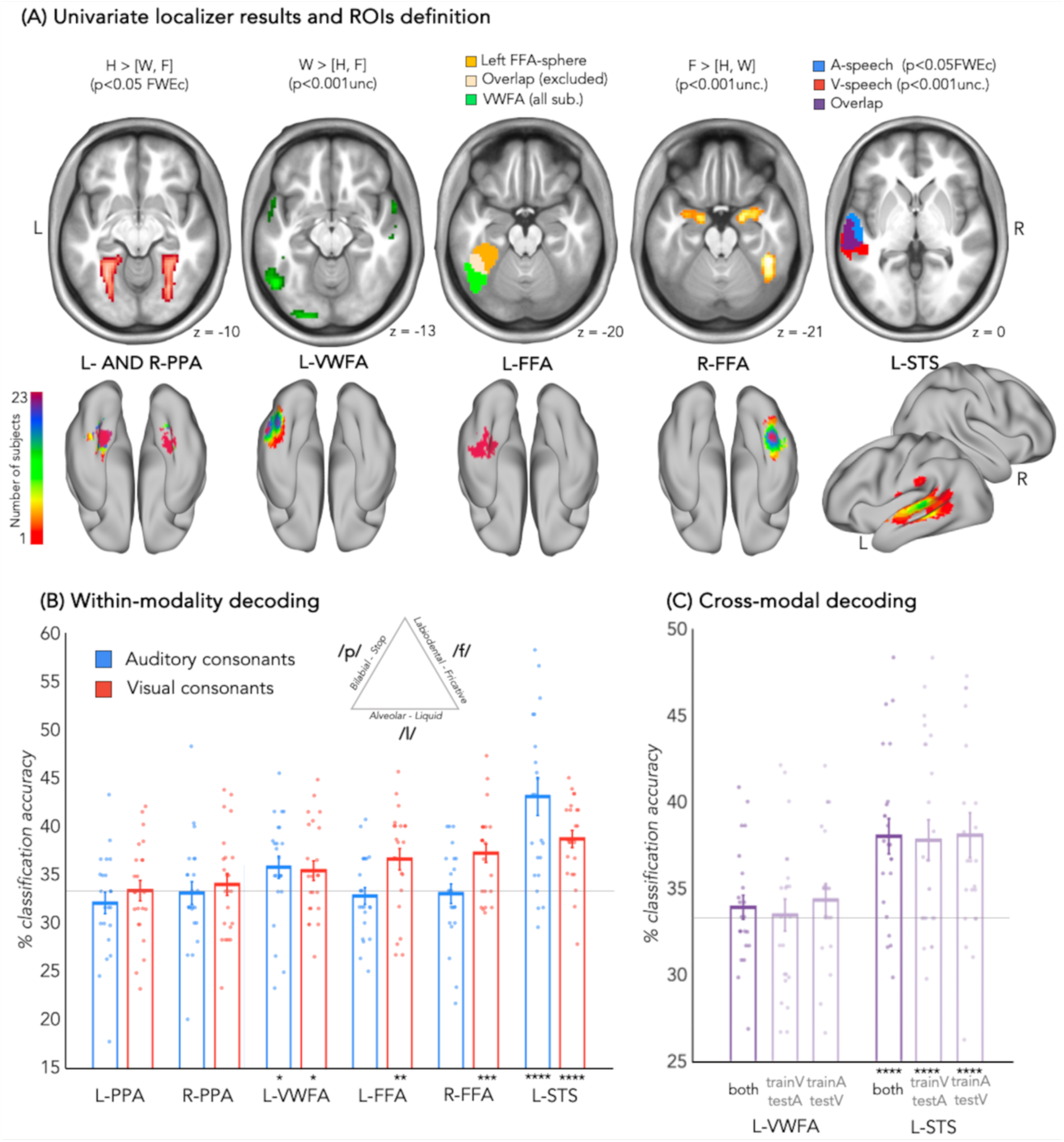
Univariate and multivariate results. (A) Four first panels represent group-level univariate contrasts to functionally define PPA (in red, house>others), VWFA (in green, words>others) and FFA (in orange, faces>others). Only right FFA could be found at the group-level and in most subjects. Left FFA mask is defined at the group-level, from a sphere of 14mm starting from the coordinate [-36; -42; -20] and excluding voxels overlapping with any individual VWFA-masks (light orange). The 5^th^ panel represents group-level left auditory speech activations in blue and visual speech activations in red, as well as voxels responding to both auditory and visual speech at the group level in purple, for visualization purpose only. The five lower panels represent the overlap and variability of each individually defined ROIs. (B) Within-modality decoding accuracies in all the individually-defined ROIs. Results are FDR-corrected for 6 ROIs (*p < 0.05, **p < 0.01, ***p < 0.001, ****p < 0.0001). Dots represent individual subjects’ classification accuracies. Error bars represent the standard error of the mean. This panel also includes a schematic representation of the distinctive articulatory traits of the selected consonants, in order to maximize visibility and recognition. (C) Cross-modal decoding accuracies for the ROIs that showed significant auditory and visual within-modality decoding, i.e., the left VWFA and left STS, for different direction of train- and test-set as well as their average. Results are FDR-corrected for 2 ROIs (*p < 0.05, **p < 0.01, ***p < 0.001, ****p < 0.0001).

**Figure 3.**
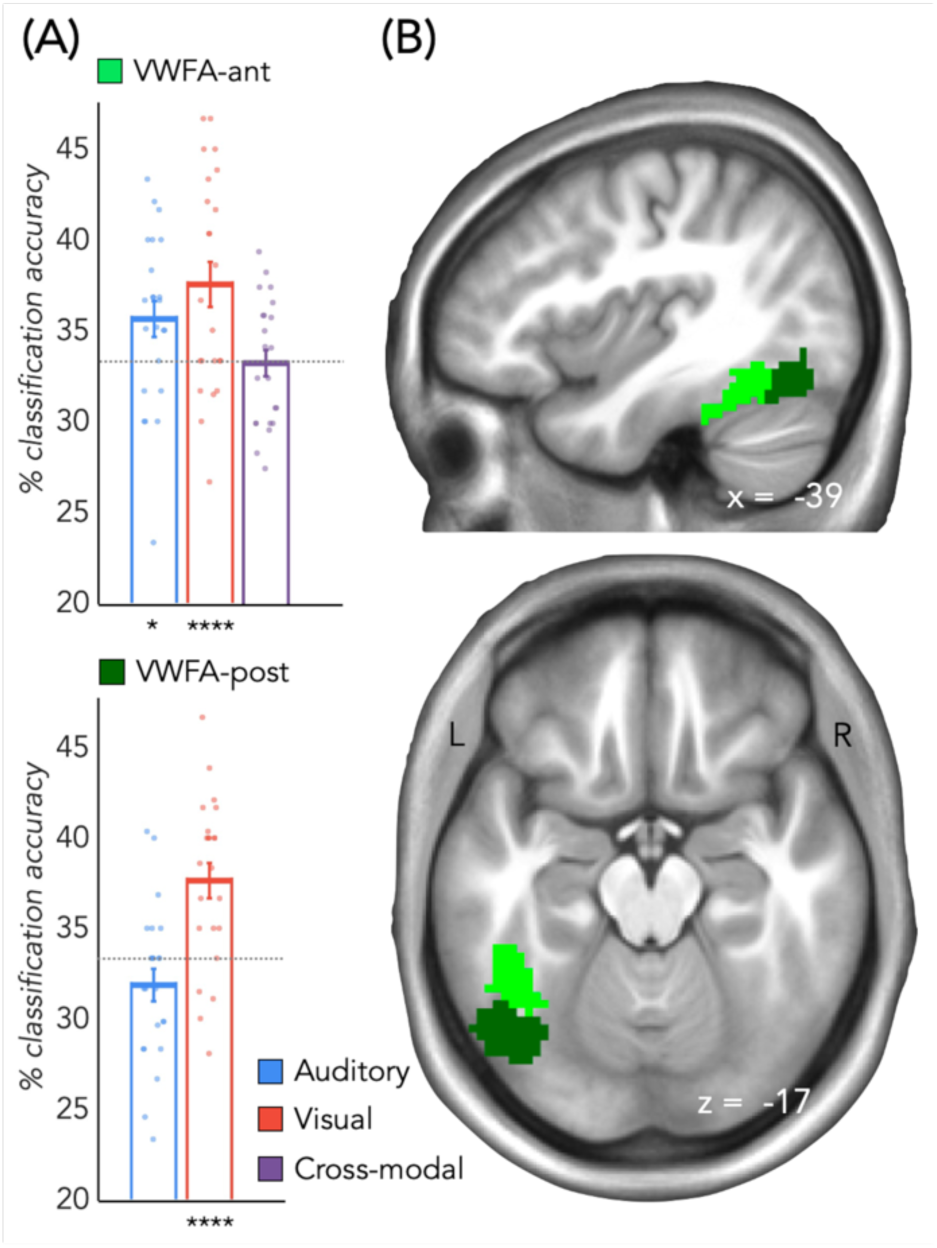
Decoding accuracies in VWFA-ant and VWFA-post. (A) Left panel represents the within- and cross-modality decoding accuracies. Results are FDR-corrected for 2 ROIs (*p < 0.05, **p < 0.01, ***p < 0.001, ****p < 0.0001). Dots represent individual subjects’ classification accuracies. Error bars represent the standard error of the mean. (B) Right panel illustrates the group-defined VWFA-ant (in light green) and VWFA-post (in dark green).

#### 2.4.5 Multisensory speech-ROI definition

We combined the results from our Auditory- and Visual-Speech localizers to identify the left area responding to speech from both modalities. In each subject, we identified voxels sensitive to auditory speech using the contrast [auditory speech] > [scrambled speech] at a threshold of p<0.05 FWE-corrected, and voxels sensitive to visual speech with the contrast [visual speech] > [non-linguistic lip movements] at a threshold of p<0.001 or p<0.01 uncorrected depending on the individual activity map. We then selected the voxels showing selectivity in the two modalities to draw an overlap cluster, focusing on the left hemisphere due to the absence of significant cluster in the right hemisphere at p<0.05 FWE-corrected for visual speech, and consistent with the left-lateralized TVSA described by Bernstein et al. (2011). The size of the overlap clusters varied considerably from one subject to another (from 6 to 329 voxels) making it impossible to use these clusters directly as ROIs in the subsequent analytical steps. Thus, we extracted visual peak coordinates of each subject in its own overlap cluster and drew an expanding ROI of 10mm around it. We did not run the Visual-Speech localizer experiment in two subjects due to time constraints. In one of the subjects who underwent the Visual-Speech localizer experiment, we did not find an overlap for auditory and visual speech. For these three subjects, we used a group-defined overlap-cluster. More precisely, we identified group-level significant activation clusters for [auditory speech] > [control condition] and [visual speech] > [control condition], at a cluster-level threshold of p<0.05 FWE-corrected and p<0.001 uncorrected, respectively. We defined the overlapping voxels between the visual and auditory maps as the group-level overlap (Fig. 2A).

#### 2.4.6 MVP decoding within modality in ROIs

We performed a multiclass (3-way) multivariate pattern (MVP) decoding using an SVM classifier to explore the representation of the consonants within each of the 6 ROIs (left VWFA, left and right FFA, left and right PPA, left TVSA), defined by the corresponding binary masks described above. Multivariate analysis was performed using CoSMoMVPA (http://www.cosmomvpa.org; implemented in MATLAB; Oosterhof et al., 2016). One beta image was estimated for each consonant per run, averaging the 9 presentations of the same consonant (3vowels*3speakers) during each run, ensuring diversity of low-level acoustic or visual features. For each modality, a linear SVM classifier was trained to discriminate the activation patterns elicited by the 3 consonants using the data of N-1 runs (training set). The performance of the SVM classifier trained on this training set was then evaluated on the remaining run (test data) in each subject individually, following a leave-one run-out cross-validation scheme. These steps were repeated as many times as we had runs for each subject (N-fold cross-validation; e.g., 20 times for the subjects who performed 20 runs), so that the test data comprised a different run in every classification fold. We used feature-selection on the training data in each cross-validation fold to use the most informative 200 voxels in each subject and each ROI and applied the same feature-selection on the test set. A single classification accuracy was then obtained by averaging the classification accuracies for all the cross-validation folds. If the classifier could successfully discriminate the different consonants above chance (33%), this would indicate the presence of phonological representation of these 3 consonants in the tested modality that the classifier was able to learn during the training. The analysis procedure was identical in the two modalities.

#### 2.4.7 MVP decoding across modality

In the regions that showed significant above chance classification of the consonants in the auditory and visual modalities, a cross-modal classification analysis was performed to test whether the representational geometry of the consonants was partially shared across modalities. Patterns of activation were demeaned individually in each modality, to equate the mean activity across auditory and visual runs. The SVM classifier was trained on one modality (e.g., vision) and tested on the other modality (e.g., audition), and reversely. We used feature-selection on the training data in each cross-validation fold to use the most informative 200 voxels in each subject and each ROI. For each subject in each ROI, we obtained 2 cross-modal decoding accuracies, one for the cross-validation folds using audition as training set and vision as test set, and one for the cross-validation folds using vision as training set and audition as test set. A two-way cross-modal decoding accuracy was finally obtained by averaging the classifications of all cross-validation folds in the 2 directions (Rezk et al., 2020).

#### 2.4.8 Statistical analyses for MVP decoding results

Significance of the decoding accuracies was evaluated through a non-parametric procedure involving permutations and bootstrapping of the data (Stelzer et al., 2013) in the following way: for each subject, one-hundred MVP-classification procedures were performed, with the labels for the consonants randomly assigned. Then, one value from the obtained one hundred decoding accuracies was sampled from each subject, and this operation was repeated ten thousand times, to obtain a group-level null distribution composed of 10.000 values. The statistical significance of the classification results was assessed by calculating the proportion of accuracy values in the null distribution that exceeded the observed value. All p-values were then FDR corrected for multiple comparisons (number of comparisons equal to the number of ROIs tested, e.g., 6 for the within-modality decoding and 2 for the cross-modal decoding analysis). P-values for the comparison between VWFA-ant and VWFA-post were also FDR corrected, with the number of comparisons set to 2.

#### 2.4.9 MVP decoding searchlight-approach

In addition to the ROI-decoding approach, we implemented a whole-brain cross-modal searchlight to detect regions showing cross-modal decoding outside of our a priori ROI space. Since this analysis was exploratory (i.e., not based on pre-defined ROI), it relies on group analyses. Consequently, it does not benefit from precise functional and individual definition and requires whole-brain statistical correction for multiple comparisons (Zhan et al., 2023). Using CoSMoMVPA, a sphere of 100-voxels was moved across the whole brain of each subject, taking one by one each voxel of the brain as the center of the searchlight sphere. Within each sphere, a cross-modal decoding was performed similarly to the paradigm described in the previous section (section 2.4.5). The individual searchlight approach generated one cross-modal decoding accuracy map for each subject after averaging the 2 directions of the cross-modal decoding (train on vision, decode on audition and reversely). Chance level accuracy (33%) was subtracted from the decoding values. These above-chance individual accuracy maps were smoothed with 8 mm FWHM smoothing kernel and entered in a second-level model. A one sample t-test was performed on the searchlight maps to test if the values were above 0 (chance level already subtracted from the data). The results were thresholded at cluster level with a Family Wise Error multiple comparisons correction (FWEc). Cluster extent was defined with initial voxel-wise threshold of *p* < 0.001, resulting in a cluster size of 297 voxels.

## 3. Results

### 3.1 Univariate localizers results

In the Word-Face-Scene localizer, the univariate contrast [words] > [faces and houses] revealed a significant cluster including 3 voxels in the VOTC at threshold of p<0.05 FWE, with a peak coordinate at [x=-52; y=-63; z=-13]. Additionally, we identified a significant cluster for the contrast [faces] > [words and houses] in the right VOTC at the same threshold, at coordinates [x=44; y=-44; z=-21] and including 23 voxels. Given the small size of the left VWFA and right FFA clusters, we used a statistical threshold of p<0.001 uncorrected for ROI definition (see Method section). No face-selective cluster was found in the left VOTC even at a lower threshold of p<0.001 uncorrected. Lastly, the contrast [houses] > [faces and words] highlighted two distinct clusters in the left and right VOTC at a threshold of p<0.05 FWE. The activation peaks were located in [x=29; y=-68; z=-10] on the right and [x=-29; y=-57; z=-10] on the left and the clusters comprised 888 (extending to occipital and parietal regions) and 301 voxels respectively (Fig 2A).

The univariate contrast of [auditory speech] > [scrambled speech] in the Auditory-Speech localizer revealed two significant clusters in the left and right superior temporal gyrus, at a cluster-level threshold of p<0.05 FWE. The right cluster comprised 355 voxels with a peak coordinate at [x=60; y=-3; z=-5]. The left cluster comprised 413 voxels with a peak coordinate at [x=-60; y=-11; z=3]. The univariate contrast of [visual speech] > [non-linguistic lip movements] in the Visual-Speech localizer revealed one cluster sensitive to visual speech more than non-linguistic lip movements corrected at a cluster-level threshold of p<0.05 FWE. This cluster was in the left posterior superior temporal sulcus with a peak coordinate at [x=-57; y=-31; z=3] and comprised 8 significant voxels. To determine the group-level audio-visual region, we used the same cluster at a lower statistical threshold of p<0.001 uncorrected, in order to include enough voxels in the overlap (Fig. 2A).

### 3.2 Behavioral results in the phonological task

D’ values were calculated individually, for the auditory and visual runs separately. The mean d’ in auditory task was 3.893 (SD = 0.478) and 2.963 (SD = 0.650) for lipreading. The results of the paired-sample t-test showed a significant difference for the auditory and visual tasks (t = 6.5586, df = 21, p<0.0001) indicating that d’ primes were higher in audition than in vision. Even in the most difficult modality (visual), the d prime values were high, with participants able to identify the repeated syllables well (mean percentage of target detection in visual modality = 75.99%; in auditory modality = 89,57%).

### 3.3 Auditory and visual decoding accuracies

We performed multiclass within-modality decoding of the 3 consonants (F, P, L) presented in visual and auditory modality, in each of the individually defined ROIs. Results of this analysis are shown in Figure 2.B. In the VWFA, we were able to decode consonants presented in the auditory modality (mean decoding accuracy, or M_DA_ = 35.9%; p_FDR_=0.0188) and in the visual modality (M_DA_ = 35.6%; p_FDR_=0.0276). No significant decoding was observed in the left and right PPA for the auditory consonants (M_DA_ = 33.1%; p_FDR_ = 0.8380 for right PPA; M_DA_ = 32.1%; p_FDR_ = 0.8790 for left PPA) nor for the visual consonants (M_DA_ = 34%; p_FDR_ = 0.2614 for right PPA; M_DA_ = 33.5%; p_FDR_ = 0.4953 for left PPA). In the right FFA, we observed above-chance classification of the visual consonants (M_DA_ = 37.3%; p_FDR_=0.0003) but not for the auditory consonants (M_DA_ = 33%; p_FDR_ = 0.8380). In the multisensory-speech ROI located in the left superior temporal sulcus (lSTS), the decoding accuracies were significantly above chance in the auditory (M_DA_ = 43.1%; p_FDR_<0.0001) and visual (M_DA_ = 38.7%; p_FDR_<0.0001) modality. To further explore the unique multisensory nature of VWFA in the fusiform cortex, we defined a left FFA ROI at the group level (see Method Section). Similar to what we observed in the right FFA, we could successfully decode the visual consonants (M_DA_ = 36.6%; p_FDR_=0.0048) but not the auditory consonants (M_DA_ = 32.8%; p_FDR_=0.8124). This suggests that the significant decoding of auditory consonants observed in the left VWFA is specific to the word-sensitive cortical patch, and not a general property of the left VOTC (see also the absence of decoding in the left PPA). Additionally, we applied the same decoding procedure in VWFA-ant and VWFA-post. We observed significant decoding accuracies in both regions in vision (M_DA_ = 37.5%; p_FDR_<0.0001 for VWFA-ant; M_DA_ = 37.6%; p_FDR_<0.0001 for VWFA-post) but only in the anterior part in audition (M_DA_ = 35.6%; p_FDR_ = 0.0286 for VWFA-ant; M_DA_ = 31.8%; p_FDR_ = 0.9219 for VWFA-post) (Fig. 3).

### 3.4 Cross-modal decoding accuracies

We performed MVP-classification across modalities in the regions that showed significant within-modality decoding accuracies in both conditions, namely the VWFA and the lSTS. Results are shown in Fig. 2C. The mean cross-modal decoding accuracy was not significantly above chance in the VWFA (M_DA_ = 33.9%; p_FDR_ = 0.2753) indicating no alignment across modalities in the phonological representation in VWFA, while it was significant in the lSTS (M_DA_ = 38%; p_FDR_<0.0001), suggesting (partially) shared patterns of representation in the auditory and visual modalities in this region. Similarly, we did not observe cross-modal decoding in VWFA when restricting our analyses to the anterior portion (M_DA_ = 33.3%; p = 0.4713; see Fig. 3).

### 3.5 Cross-modal whole-brain searchlight

To further explore the presence of multisensory phonological networks beyond our ROIs, we relied on a whole-brain cross-modal searchlight approach (Kriegeskorte et al., 2006). Two significant clusters could be identified at a threshold of p<0.05 FWEc corrected (Fig. 4). The first cluster comprised 297 voxels and was located in the right posterior superior temporal sulcus (right pSTS; peak coordinates at x=60; y=-24; z=0). The second cluster comprised 749 voxels and encompassed several significant peaks and anatomical areas. One peak was situated in the left pSTS, confirming the results obtained in the ROI-approach (peak coordinates at x=-57; y=-34; z=5). Finally, two peaks were significant in the left central opercular cortex and the ventral part of the left pre/post-central gyrus (peak coordinates at x=-55; y=-13; z=18 and x=-52; y=-13; z=29, respectively). In order to test the overlap of this somato-motor region with oro-facial articulators involved in speech production, we performed a small volume correction (SVC) using a 10mm radius sphere centered on coordinates of the literature. The SVC was applied on a motor region involved during speech production (MNI coordinates [x=-61; y=0; z=16] from Kern et al., 2019) and on a region selective for non-speech oro-facial movements (MNI coordinates [x=-56; y=-4; z=32] from Grabsky et al., 2012). Both SVCs gave significant results (p<0.05 FWE-corrected), indicating a specific involvement of oro-facial sensori-motor cortex for aligned multisensory representation of visemes and phonemes.

**Figure 4.**
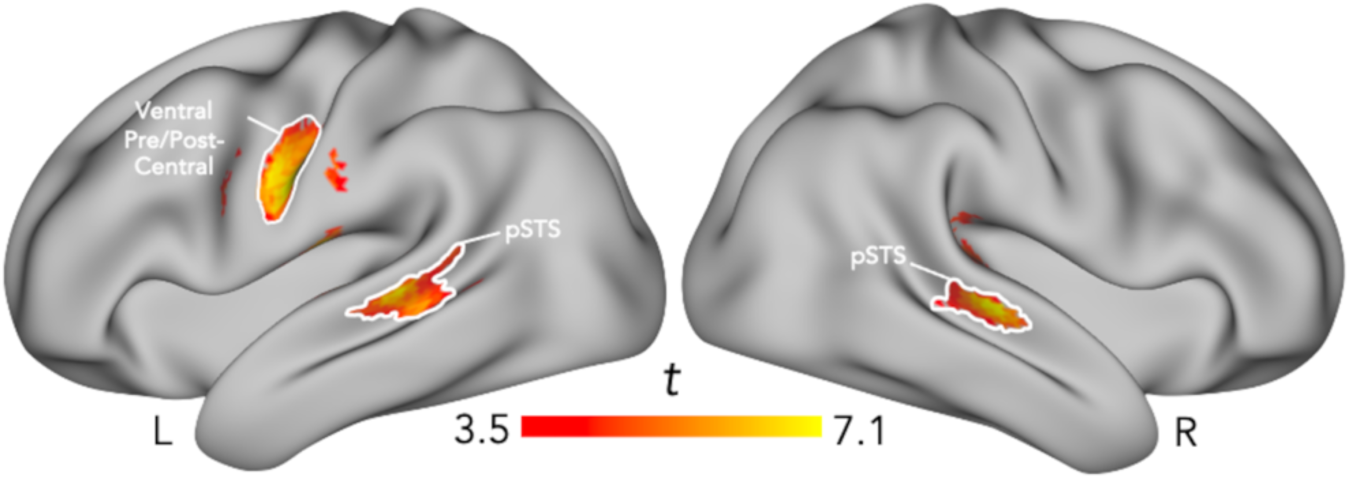
Cross-modal whole-brain searchlight decoding. Statistical map of the cross-modal whole-brain searchlight, displayed at a statistical threshold of p<0.05 FWE-cluster corrected. The three anatomical clusters are delineated.

## 4. Discussion

We examined auditory, visual, and multisensory phonological representations using uni- and cross-modal fMRI MVP-decoding analyses within individually and functionally defined brain regions of the VOTC and STS, complemented by a whole brain cross-modal decoding approach.

One of the most striking findings of our study is the observation of significant within-modality decoding of consonants in both visual and auditory modalities in the VWFA. When distinguishing between anterior and posterior parts of the VWFA, we observed significant decoding in both vision and audition only in the anterior part, corroborating previous evidence of a posterior-to-anterior gradient of visual-to-linguistic processing in the word-selective cortex (Lerma-Usabiaga et al., 2018). Importantly, neither the posterior VWFA nor ipsilateral FFA and PPA showed significant decoding in the auditory modality, indicating that the presence of multisensory phonological representations in the left VOTC is specific to the VWFA. Our results support recent magneto- and electro-encephalography (M/EEG) studies showing neural tracking of visuo-phonological units (visemes) (O’Sullivan et al., 2017; Bröhl et al., 2022; Nidiffer et al., 2023) and of unheard auditory speech signal (Hauswald et al., 2018) in the occipital cortex during silent lipreading. These findings suggested that some visual regions support auditory and visual linguistic representations, but M/EEG’s low spatial resolution prevented the identification of functionally specific visual areas. Our results reveal the specific role of the anterior VWFA in multisensory phonological computations during speech processing. Interestingly, despite VWFA representing phonemes and visemes, our data showed no significant cross-modal decoding in this region, indicating a lack of shared representational format.

What is therefore represented in VWFA? One might argue for an automatic co-activation of orthographic code during auditory and/or visual speech perception (Dehaene et al., 2010; Hauw et al., 2023; but see Pattamadilok et al., 2007), leading to significant decoding in VWFA. However, significant cross-modal decoding would be expected if the written code were automatically activated during auditory and visual speech processing, as this format would be shared regardless of the modality. Consistent with the idea that phonological representation beyond orthography can be observed in VWFA, recent research indicates that the VWFA’s sensitivity to auditory speech and phonological processing is evident in preliterate children (Li et al., 2023), suggesting its predisposition towards spoken material before developing visual selectivity for written words. Secondly, auditory phonological representations in the VWFA could be attributed to visual imagery (i.e., imagining facial patterns during the auditory task). However, if true, significant auditory-decoding would also be expected in the FFA, which show significant decoding of visemes and is involved in visual imagery of facial patterns as well (O’Craven and Kanwisher, 2000). Thus, visual imagery and automatic co-activation of written code are unlikely to fully explain the multisensory phonological representations observed in the VWFA. The absence of aligned representation across visemes and phonemes found in this study is coherent with recent findings from a transcranial magnetic stimulation (TMS) adaptation study revealing facilitatory effects of TMS applied on the VWFA during a lexical decision task with written or spoken words, but only within the same modality as the adaptation phase (Pattamadilok et al., 2019). Within-modality adaptation for both auditory and visual stimuli, coupled with the lack of cross-modal adaptation, suggests that the VWFA contains segregated auditory and visual neural populations coding phonological representations.

Our results challenge the traditional view of the VWFA as a purely visuo-orthographic system (Dehaene et al., 2005). Rather, it seems that the VWFA is involved in visual, auditory and audio-visual language processing, such as reading, lipreading and speech processing, supporting an integrative role in the language system (Price and Devlin, 2011; Qin et al., 2021). Some authors suggested that the VWFA’s neural tuning for orthographic units, or graphemes, may result from recycling its ability to process visual linguistic information like lip movements, or visemes (Bernstein et al., 2014; Hannagan et al., 2015). The VWFA’s connectivity with superior temporal auditory regions (Stevens et al., 2017; Saygin et al., 2016), essential for grapho-phonological conversion in reading, may also facilitate audio-visual matching during speech perception. Further studies should characterize VWFA’s connectivity with other sensory and associative areas during reading and lipreading to explore processing similarities between these two visual forms of language.

We successfully decoded visemes but not phonemes in the right and left FFA, suggesting that the FFA encodes categories of lip movements or facial configurations, contrary to the classical idea that FFA only encodes invariant aspects of faces like identity (Haxby et al., 2000). Our results do not clarify whether FFA encoded our visual stimuli as dynamic (lip movements) or static (lip configuration), nor whether these computations are language-specific or related to facial pattern recognition independently of linguistics. Our results however align with recent evidence showing that FFA also encodes dynamic facial patterns of emotions (Ganel et al., 2005; Mattioni, Falagiarda et al., 2024), suggesting FFA may involve in recognizing communicative facial patterns more generally, and is not specific to language.

Confirming previous results (Bernstein et al., 2011), we identified a left pSTS region sensitive to visual speech through our Visual-Speech localizer. By combining these visual univariate results with the Auditory-Speech localizer activations, we delineated a “multisensory speech region” in each subject’s lSTS, activated by both auditory and visual speech. Our results showed auditory and visual phonological representations in the lSTS and significant cross-modal decoding of phonological units, suggesting aligned phonological activations across modalities. A complementary cross-modal searchlight confirmed significant cross-modal decoding not only in the lSTS but also in its right counterpart (see Fig. 4). The presence of cross-modal decoding in STS indicates that phonological representations, known for auditory speech (Chang et al., 2010; Bonte et al., 2014), are also accessible from vision and shared across the senses in a partially abstracted format. This aligns with findings of high-order encoding of speech sounds in the language cortex (Chang et al., 2010; Leonard et al., 2023), and shows phonological representation independent of spectrotemporal acoustic cues (Mesgarani et al., 2014). Recent intracranial encephalography (iEEG) studies in pSTG demonstrated that audio-visual activations were weaker than auditory-only activations (Karas et al., 2019; Metzger et al., 2020). The authors concluded that perceived visuo-phonological features modulate auditory cortex activity by suppressing responses to incompatible auditory phonemes (i.e., subadditive responses). Our findings may support this hypothesis, as the multivariate classification approach relates to activation patterns independently of response amplitude. Thus, activation and de-activation patterns elicited by visemes and phonemes may lead to significant cross-modal decoding, indicating a shared representation accessible from both modalities.

The whole-brain cross-modal decoding approach also revealed cross-modal activations in the left ventral somato-motor cortex, corresponding to regions known to represent oro-facial articulators such as the lips, jaw and tongue involved in speech production (Grabsky et al., 2012; Kern et al., 2019). One possibility is that both auditory (Lankinen et al., 2023) and visual speech perception co-activate somato-motor patterns (Watkins et al., 2003). Alternatively, the somato-motor cortex might support an abstract phonological representation as suggested by an fMRI study using representational similarity analysis (RSA) and showing that speech representations in the somato-motor cortex are independent of acoustic features (Evans and Davis, 2015). Finally, findings from intracranial EEG suggested that the motor cortex supports both acoustic and articulatory patterns (Cheung et al., 2016). Supporting the causal role of somato-motor cortex in speech representation, facial skin deformation disrupts speech perception (Ito et al., 2009) and TMS-pulses on the lip or tongue motor cortex facilitates word perception (Schomers et al., 2015). It would be interesting to test whether such perturbations similarly affect visual and auditory speech perception to assess the somato-motor cortex’s causal role in representing phonology from both modalities.

Altogether, our results support an integrative model where multisensory phonological representations are distributed across several sensory and functional networks, including the auditory, visual and somato-motor cortices (Schomers and Pulvermüller, 2016). More specifically, the involvement of the VWFA (and in particular its anterior portion) in representing co-localized but not aligned auditory and visual phonetic information supports a linguistic function of this region beyond orthographic coding (Price and Devlin, 2011; Pattamadilok et al., 2019; Wang et al., 2022), acting as an integrative relay between the visual and audio-motor system, where visemes and phonemes merge into a partially aligned/abstracted representation.

## Acknowledgements

The project was funded in parts by a Mandat d’Impulsion Scientifique awarded to OC, the Belgian Excellence of Science (EOS) program (Project No. 30991544) awarded to OC, and a Flagship ERA-NET grant SoundSight (FRS-FNRS PINT-MULTI R.8008.19) awarded to OC. AV is a research fellow and OC is a senior research associate at the Fond National de la Recherche Scientifique de Belgique (FRS-FNRS).

## Data and code availability statement

Codes and stimuli used to perform fMRI experiments (decoding and localizer experiments) are available on GitHub for the scripts (https://github.com/avanaudenhaege/LipSpeech_project) and OSF for the stimuli (https://osf.io/2xtsn/). Codes for analyses of the data is available on GitHub (https://github.com/avanaudenhaege/LipSpeech_analyses). Raw defaced MRI data are available on GIN in BIDS format (https://gin.g-node.org/avanaudenhae/LipSpeech-RAW).

## Author contributions

See Figure 5.

**Figure 5.**
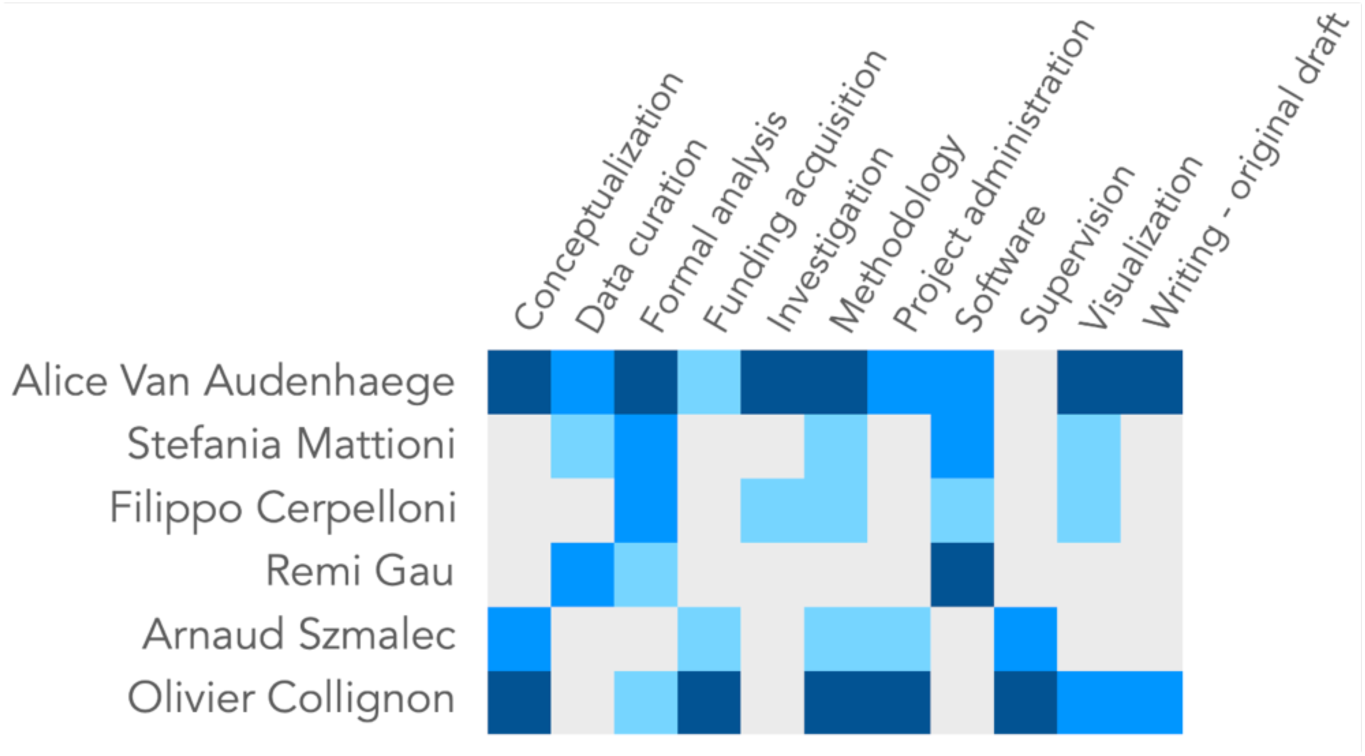
Author contributions. Based on the CRedIT (https://credit.niso.org/) taxonomy. Each contribution was assessed as ‘lead’ (dark blue), ‘significant contribution’ (mid blue), ‘support’ (light blue).

